# miR-203 controls developmental timing and early fate restriction during preimplantation embryogenesis

**DOI:** 10.1101/2024.02.06.579214

**Authors:** José González-Martínez, Agustín Sánchez-Belmonte, Estefanía Ayala, Alejandro García, Enrique Nogueira, Jaime Muñoz, Anna Melati, Daniel Giménez, Ana Losada, Sagrario Ortega, Marcos Malumbres

**Affiliations:** Cell Division and Cancer group, Spanish National Cancer Research Centre (CNIO) Madrid.; Cancer Cell Cycle group, Vall d’Hebron Institute of Oncology (VHIO), Vall d’Hebron Barcelona Hospital Campus, Barcelona, Spain; Mouse Genome Editing Unit, Spanish National Cancer Research Centre (CNIO) Madrid.; Chromosome Dynamics group, Spanish National Cancer Research Centre (CNIO) Madrid.; ICREA, Barcelona

**Keywords:** Embryo development, Developmental timing, Histone acetylation, microRNA, Pluripotency, Totipotency

## Abstract

Commonly expressed at developmental transitions, microRNAs operate as fine tuners of gene expression to facilitate cell fate acquisition and lineage segregation. Nevertheless, how they might regulate the earliest developmental transitions in early mammalian embryogenesis remains obscure. Here, in a strictly in vivo approach based on novel genetically-engineered mouse models and single-cell RNA sequencing, we identify miR-203 as a critical regulator of timing and cell fate restriction within the totipotency to pluripotency transition in mouse embryos. Genetically engineered mouse models show that loss of miR-203 slows down developmental timing during preimplantation leading to the accumulation of embryos with high expression of totipotency-associated markers, including MERVL endogenous retroviral elements. A new embryonic reporter (eE-Reporter) transgenic mouse carrying MERVL-Tomato and Sox2-GFP transgenes showed that lack of miR-203 leads to sustained expression of MERVL and reduced Sox2 expression in preimplantation developmental stages. A combination of single-cell transcriptional studies and epigenetic analyses identified the central coactivator and histone acetyltransferase P300 as a major miR-203 target at the totipotency to pluripotency transition in vivo. By fine tuning P300 levels, miR-203 carves the epigenetic rewiring process needed for this developmental transition, allowing a timely and correctly paced development.

## Introduction

The fundamental principles governing the acquisition of cell lineage diversity from a totipotent zygote to a multicellular organism during embryogenesis have fascinated scientists for decades. Preimplantation embryonic development is a meticulously orchestrated process characterized by tightly regulated cell transitions, each marking distinct stages of cellular specialization. In this process, the totipotent zygote harboring full developmental potential undergoes several fate-restricting developmental transitions until the formation of the blastocyst, which will then implant in the uterine wall (Menchero *et al*, 2017; Wilkinson *et al*, 2023). Importantly, control of developmental timing is known to be essential for this process.

Developmental timing is known to rely on epigenetic modifications (Ciceri *et al*, 2024), which dynamically regulate the chromatin landscape and gene expression patterns contributing to the establishment of cellular identity, especially during developmental transitions (Wilkinson *et al*., 2023). A critical developmental transition in early embryogenesis is the 2-cell-to-8-cell transition, in which lineage segregation and fate restriction occur when repressive H3K27 tri-methylation (H3K27me3) marks are progressively replaced by H3K27 acetylation activatory marks (H3K27ac) in the enhancers and promoters of genes transcriptionally induced (Gao *et al*, 2018; Lu *et al*, 2021; Millan-Zambrano *et al*, 2022). How this switch occurs in different species and cell lineages is an outstanding question with critical implications in early embryonic development (Wilkinson *et al*., 2023). However, the scarcity of in vivo studies addressing these issues makes the identification of these core developmental regulators still elusive. miRNAs were originally identified as regulators of the timing of developmental transitions during body patterning in *C. elegans* (Lee *et al*, 1993; Reinhart *et al*, 2000) and constitute attractive candidates as molecular regulators of early developmental transitions in mammalian embryogenesis. However, the role of specific miRNAs in early embryogenesis remains majorly unexplored due to their high functional redundancy and as they are opaque to most sequencing methods, making their identification elusive. We recently reported that a specific microRNA with a unique seed sequence, miR-203, can trigger strong hypomethylation in pluripotent cells (Salazar-Roa *et al*, 2020). miR-203 was initially identified as a microRNA with critical roles in skin differentiation (Yi *et al*, 2008) and soon recognized as a target of multiple alterations in cancer (Bueno *et al*, 2008). Ectopic expression of this microRNA leads general hypomethylation of the genome in embryonic stem (ES) cells and induced pluripotent stem (iPS) cells, in a DNMT3A/B-dependent manner, leading to a “tabula rasa” pluripotent cellular state in which cell differentiation towards multiple lineages is enhanced (Salazar-Roa *et al*., 2020). However, whether miR-203 plays a role during early embryonic development remains unexplored. In this work, we unveil the role of miR-203 as a regulator of epigenetic rewiring during the totipotency to pluripotency transition. Altogether, data provided here highlight miR-203 as a novel regulator of developmental timing and fate restriction in early mammalian embryogenesis.

## Results

### miR-203 is required for timely transitions during early embryonic development

In mouse embryos, miR-203 is significantly induced at the 8-cell and morula stages, in parallel to compaction and polarization of blastomeres, and its expression levels decrease before the first cell-fate decision to generate outer-residing polar cells of the trophectoderm (TE) and apolar cells in the inner cell mass (ICM; Figure 1a). miR-203 is also undetectable in cultured ESCs derived from the ICM (Supplementary Figure 1a).

**Figure 1.**
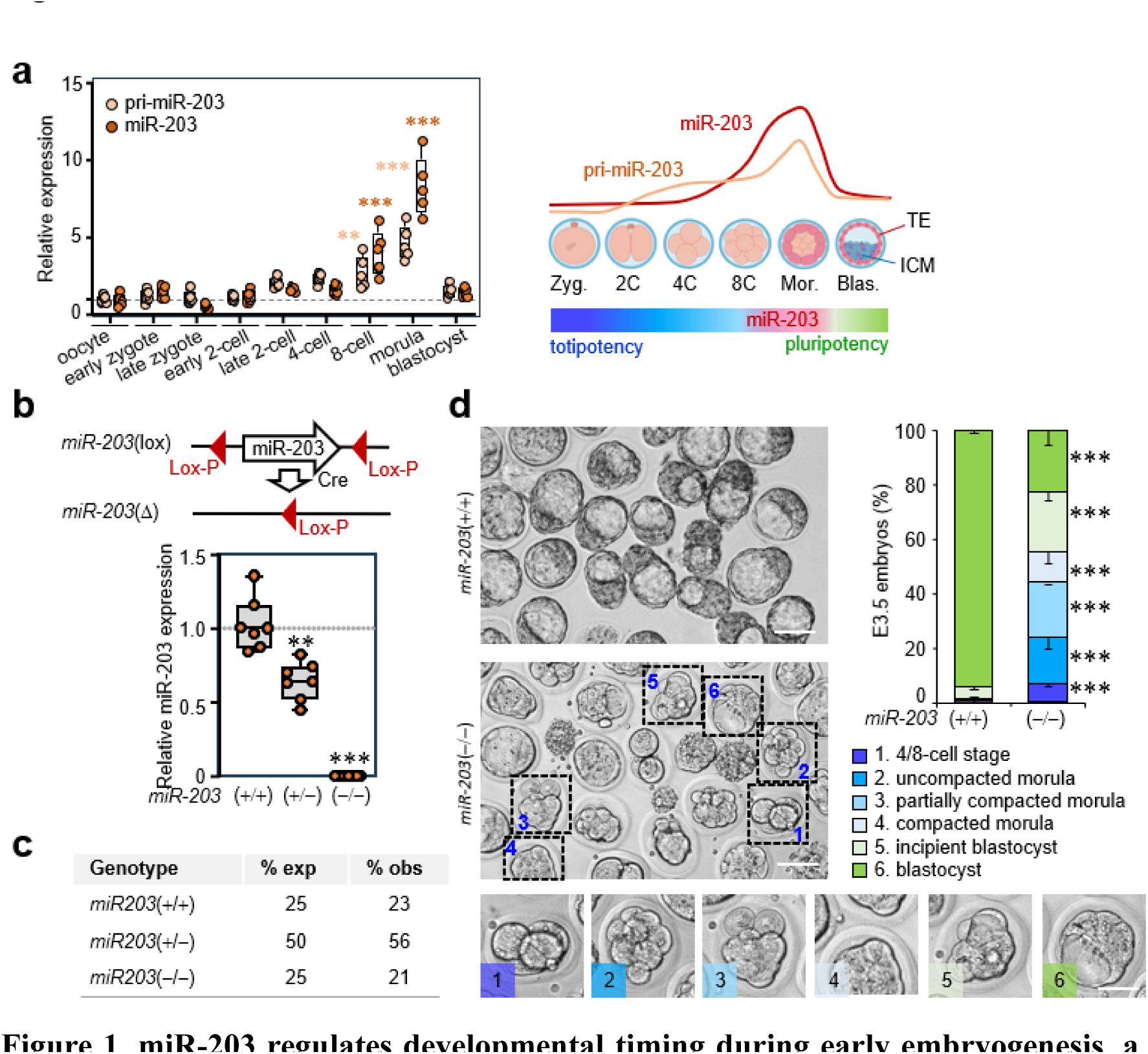
miR-203 regulates developmental timing during early embryogenesis. **a**, Expression of *pri-mmu-mir-203* (pale brown) and *mmu-mir-203* (brown) at the indicated stages during early embryonic development in the mouse. Each data point represents 15-20 embryos of the stated developmental stage from the same female donor. (n=⁓100 total embryos/experimental group). RNA expression is normalized by a housekeeping miRNA or pri-miRNA (miR-16 or pri-miR-16) that is maintained invariable during early embryogenesis. The panel to the right shows a summary of pri-miR-203 and miR-203 expression during the indicated developmental stages. **b,** Strategy used to generate miR-203-deficient mice for this work (top; see Suppl. Figure 1b for details), and relative levels of miR-203 respect miR-16 as housekeeping control miRNA in wild-type, heterozygous and homozygous null 8-cell stage embryos (20 embryos from the same female per data point). **c**, Quantification of the ratio of adult mice with the indicated genotypes in intercrosses between *miR-203*(*+*/*−*) mice. **d**. Representative images and quantification of developmental stages of *miR-203*(*+*/*+*) or *miR-203*(*−*/*−*) embryos extracted from superovulated females at E3.5. n= 84 embryos miR-203 *miR-203*(*+*/*+*) and n= 169 embryos *miR-203*(*−*/*−*). Scale bars, 100 *μ*m (50 *μ*m in the bottom insets). Data in a, b, are analyzed by a 1-way ANOVA with Tukey multiple comparisons post-hot test, and data in d are analyzed by a student t-test with Welsh correction. **, *P*<0.01; ***, *P*<0.001 of each experimental group respect early zygote in a, *miR-203*(*+*/*+*) in b and d.

To study the functional relevance of miR-203 in early developmental stages, we generated mice with a conditional deletion of the entire *mmu-mir-203* locus using the Cre-loxP system. The miR-203-encoding sequence is located at Chr. 12 in the mouse, in an intergenic region ⁓3.5-kb telomeric to *Aspg* and ⁓15-kb centromeric to the *Kif26a* gene. We placed two *loxP* sites flanking *mmu-mir-203* generating the *miR-203*(lox) allele after recombination of these sequences in ES cells (Figure 1b and Supplementary Figure 1b). Cre-mediated recombination resulted in a null allele [*miR-203*(*−*)], which was used to generate germline miR-203-deficient mice. Expression of miR-203 was reduced in *miR-203*(*+*/*−*) 8-cell-stage embryos and was undetectable in *miR-203*(*−*/*−*) homozygous mutant embryos (Figure 1b). Lack of miR-203 did not result in developmental defects as later stages as we observed no difference in the ratio of miR-203-null pups (Figure 1c), and adult *miR-203*(*−*/*−*) mice were viable with no obvious alterations apart from reduced thickness and complexity of the stratified epithelium in the skin (Supplementary Figure 1c), in agreement with a previously described role of miR-203 in skin differentiation (Yi *et al*., 2008). However, careful examination of *miR-203*(*−*/*−*) preimplantation embryos showed that lack of miR-203 resulted in a globally protracted developmental timing and kinetics of transitions in the early embryo. When analyzing the structure of E3.5 embryos directly isolated from the uterus, we observed that ⁓95% of E3.5 wild-type embryos had already reached the blastocyst stage. However, miR-203-deficient embryos displayed a significantly reduced representation of these structures (21%) in favor of more incipient blastocysts, abundant morulas at different stages as well as a few earlier (4-, 8-cell stages) embryos and a few aberrant structures (Figure 1d and Supplementary Figure 1d).

To generate detailed in vivo data about early transitions in the mouse embryo, we monitored the expression of the murine class III ERV with leucine transfer RNA primer binding site (MERVL), a set of endogenous retroviral element repeats especially expressed during the zygotic genome activation at the 2C stage which interact with totipotency-associated genes to allow their expression (Macfarlan *et al*, 2012). We generated a new transgenic mouse in which a MERVL::tdTomato reporter (Macfarlan *et al*., 2012) was injected in mouse zygotes. After selection of mice with stable germline transmission of the allele, we observed strong expression of the Tomato reporter at the 2-cell stage and decreasing expression in the subsequent developmental stages with residual or patched expression in some 8-cell embryos (Figure 2a-c and Supplementary Figure 2a). By E2.75, most *miR-203*(*−*/*−*); MERVL-Tomato 8-cell embryos displayed higher levels of expression of the Tomato reporter when compared to control embryos (Figure 2d,e and Supplementary Figure 2b). We also combined this allele with a previously generated Sox2-GFP transgenic (Suh *et al*, 2007), in which the green fluorescence reporter is strongly expressed later at the morula and blastocyst stages concomitant with the establishment of pluripotency transcriptional programs, thus generating a double MERVL-Tomato; Sox2-GFP transgenic line (early Embryo reporter or eE-Reporter mouse; Figure 2a-c). Introducing these two alleles into the miR-203 deficient mouse allowed the identification of compacted morulas with either low or high expression of MERVL-Tomato that were not present in E3.5 control embryos (Figure 2f,g), as well as morulas with separated green and red clusters of cells, suggesting not only developmental delay but also an asynchronous cell fate transition within the blastomeres of some embryos.

**Figure 2.**
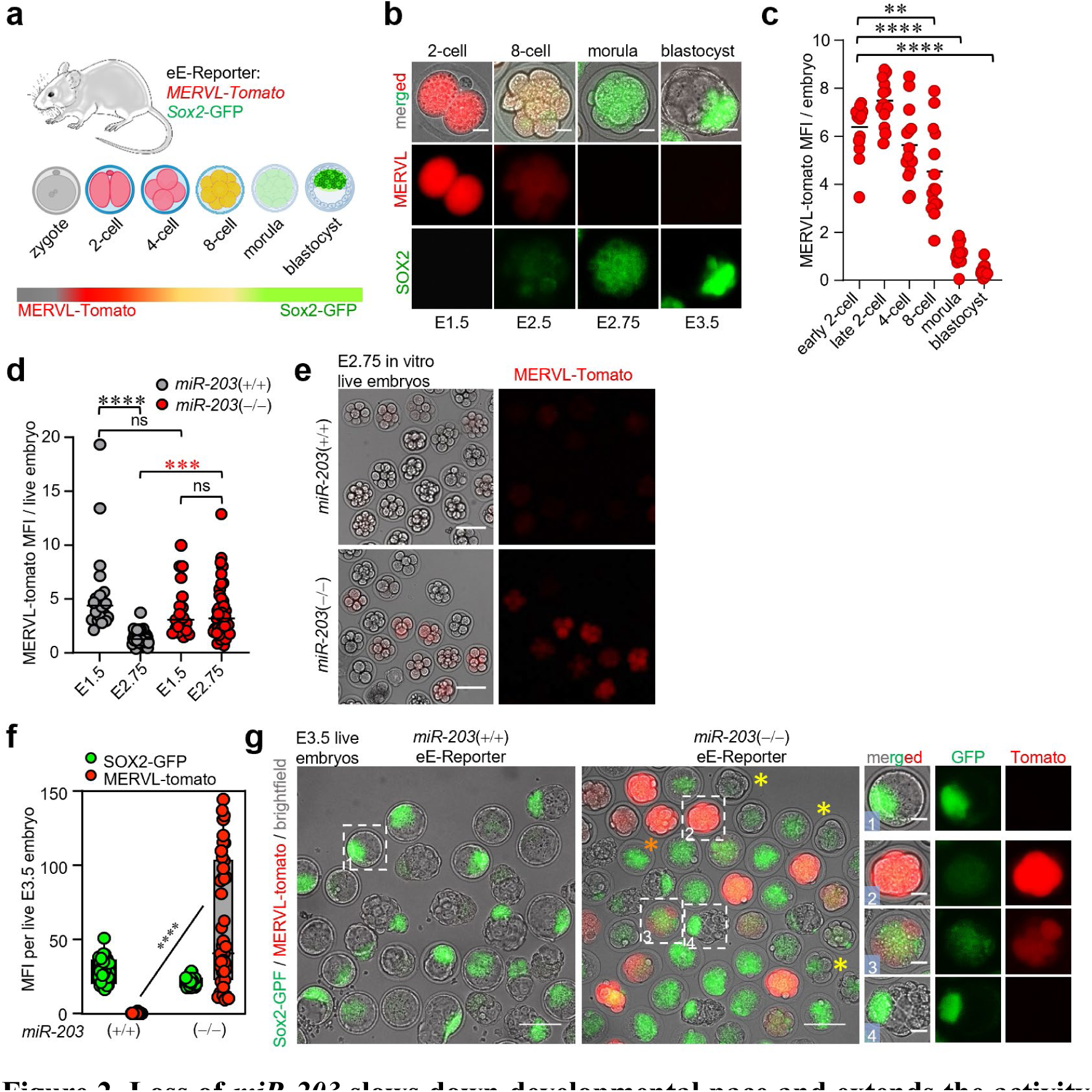
Loss of *miR-203* slows down developmental pace and extends the activity of totipotency-associated endogenous retroviral elements in vivo. **a**, Schematic representation of the eE-reporter mouse. **b**, Totipotent embryos with active MERVL retroviral elements display a peak of red fluorescence (td-tomato) in the 2-cell stage concomitant with the expression of totipotency genes, which decreases upon after the exit of totipotency in the 4-cell and 8-cell stages. Entry in pluripotency is reported by the expression of GFP under the control of the *Sox2* gene promoter at the morula and blastocyst stages. Scale bars: 20µm. **c**, Quantification of MERVL-tomato mean fluorescence intensity (MFI) in arbitrary units, per embryo, in the indicated embryonic preimplantation stages. Each data point corresponds to one embryo. Horizontal bars represent the mean of each biological group. **d**, Quantification of MERVL-tomato MFI in embryos of the indicated genotypes during E1.5 and E2.75 stages. Horizontal bars represent the mean of each biological group. Note the sustained MERVL-tomato MFI during E2.75 stage in *miR-203* deficient morulae respect wildtype controls (red asterisks). **e**, Fluorescence micrographs of live E3.5 embryos harboring the eE-Reporter and *miR-203* alleles. Insets to the right represent a *miR-203*(*+*/*+*) blastocyst bearing GFP fluorescence in the ICM (1), and *miR-203*(*-/-*) embryos with intense td-tomato fluorescence and GFP absence (2), mixed td-tomato and GFP expression (3), and GFP expression only (4). Yellow asterisks indicate embryos in the morula stage which already lost MERVL-tomato expression. All embryos harbor the *MERVL-tomato* and Sox2-GFP alleles in the same allelic dose. Scale bars: 100µm (micrographs) and 25µm (insets). **f**, Quantification of green (*Sox-GFP*) and red (*MERVL-tomato*) mean fluorescence intensity in arbitrary units (MFI) of the embryos displayed in **d**. Each data point corresponds to one embryo. **g**, Live fluorescence micrographs of the E2.75 morulas of the indicated genotypes. Scale bars: 100µm. Data in **c**, **d**, **e** are analyzed by a 1-way ANOVA with Tukey multiple comparisons post-hot test. **, *P*<0.01; ***, *P*<0.001, ****, *P*<0.0001, ns: non-significant within the indicated comparisons.

### miR-203 modulates totipotency and pluripotency programs in preimplantation embryos

To understand the molecular basis behind these defects, we collected 289 E3.5 and 540 E4.5 *miR-203*(*−*/*−*) (KO) embryonic cells, and compared them to 306 E3.5 and 1,314 E4.5 wild-type embryonic cells using single-cell RNA sequencing (scRNAseq; Supplementary Figure 3a). We also made use of an inducible knock-in allele, *miR-203*(tet), previously generated in our laboratory (Salazar-Roa *et al*., 2020), in which miR-203 can be induced by doxycycline (Dox) in the presence of the reverse transactivator rtTA (Rosa26-rtTA; Supplementary Figure 1e). Treatment of *miR-203*(+/tet); Rosa26-rtTA [from now on, *miR-203*(+/tet); with Dox in vivo resulted in a significant upregulation of miR-203 at different developmental stages (Supplementary Figure 1f). To compare with miR-203-deficient embryos, we also collected 251 E3.5 and 951 E4.5 *miR-203*(+/tet) cells after treatment with Dox when plug was detected (from now on, *miR-203*(+/tet)^Dox^). Dimensional reduction of expression data clearly separated miR-203-deficient cells from control of *miR-203*(+/tet)^Dox^ embryonic cells (Figure 3a and Supplementary Figure 3a). Initial mapping with markers of trophoectoderm (TE; *Cdx2*, *Gata3* and *Krt8*), inner cell mass (ICM; *Nanog*, *Klf4* and *Pouf5f1*) and primitive endoderm (PrE, also known as hypoblast; *Gata6, Gata4, Sox17*) suggested that a clear separation of these three lineages in E4.5 cells as well as control and *miR-203*(+/tet)^Dox^ E3.5 embryos (Figure 3b). E3.5 miR-203-deficient cells, however, were not well defined with these markers, suggesting a defect in cell fate specification.

**Figure 3.**
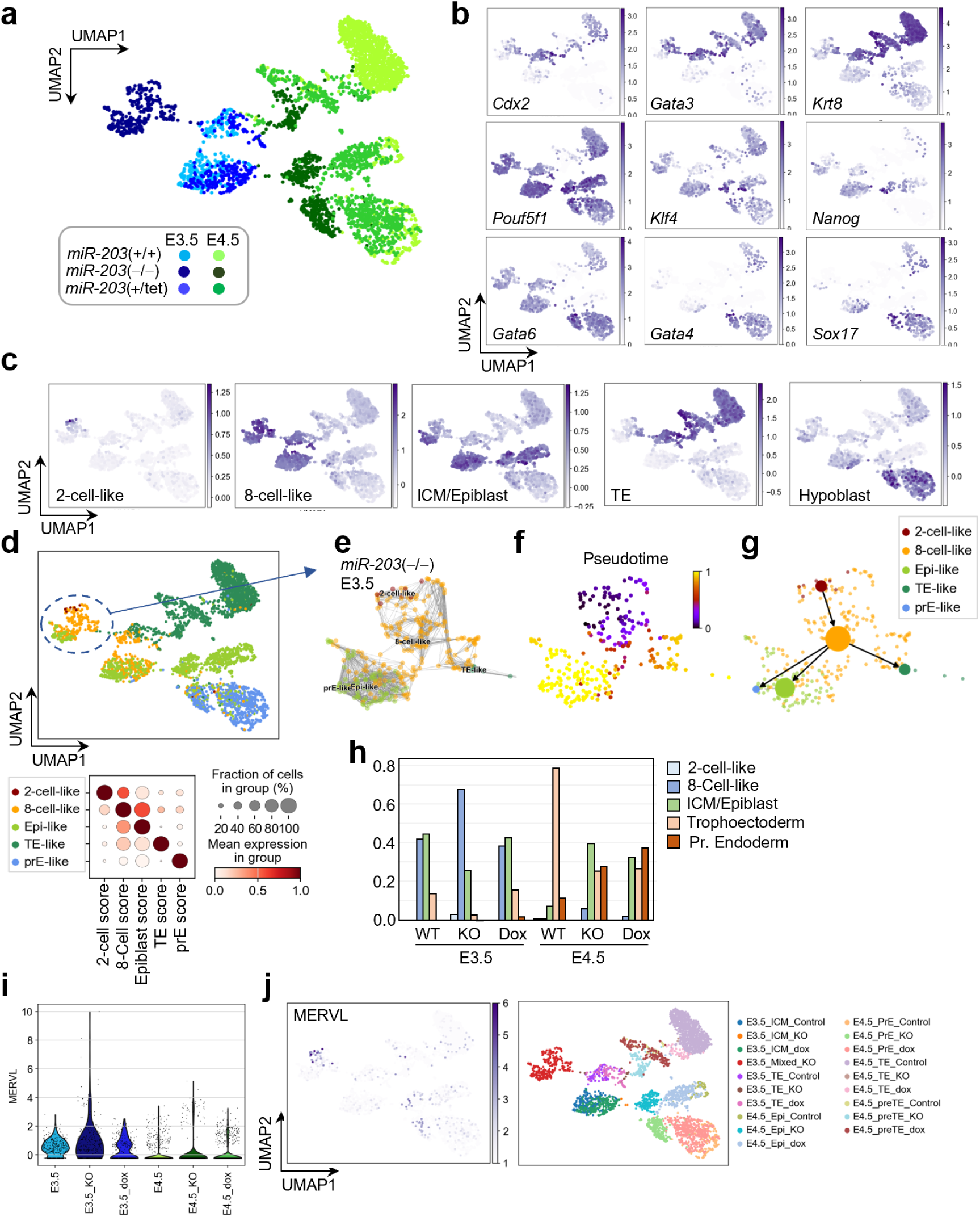
Single cell transcriptomics reveals a heterogeneous cluster of cells with protacted developmental timing and totipotency-associated transcriptional features upon genetic ablation of *miR-203 in vivo*. **a**, scRNA-seq analysis of *miR-203* wildtype, knockout and knockin embryos at E3.5 and E4.5. The UMAP plot displays six main clusters displayed below in different colors. **b**, UMAP plots displaying the expression of prototypical trophectoderm identity genes (*Cdx2*, *Gata3*, *Krt8*), inner cell mass / epiblast (*Pou5f1*, *Klf4*, *Nanog*), and primitive endoderm identity genes (*Gata6*, *Gata4*, *Sox17*). Scale bars indicate gene expression. **c**, UMAP plots showing the expression of distinctive gene signatures for 2-cell-like, 8-cell-like, inner cell mass (ICM) / epiblast, throphectoderm (TE) and hypoblast identity transcriptional programs. Scale bars indicate the gene expression. **d**, UMAP plot representing the distinct cell populations presented in **a** classified according to the lineage identity gene signatures stated in **c**. Note the presence of a hybrid, heterogeneous cluster constituted by *miR-203*(*-/-*) cells exhibiting 2-cell, 8-cell and ICM transcriptional features which cluster together (dashed circle). The dot plot below represents the numeric score for each one of the signatures in each cell cluster. **e**, Detailed UMAP plot of the *miR-203*(*-/-*) E3.5 cell cluster showing the distribution of each cell type according to the expression of the indicated lineage identity gene signatures. **f**, Pseudotime analysis projected on the *miR-203*(*-/-*) E3.5 cell cluster. **g**, PAGA velocity analysis delineating the main developmental trajectories within the *miR-203*(*-/-*) E3.5 cell cluster. **h**, Bar plot representing gene expression for each lineage identity gene signature stated in the different cell clusters. **i**, Violin plot depicting MERVL expression across the different cell clusters displayed in **a**. Note the increased expression of MERVL in *miR-203*(*-/-*) E3.5 embryos. **j**, UMAP plots showing the expression MERVL retroelements projected to the UMAP plot (left) and the separation of cells by stage, genotype and cell lineage (right).

We then used gene signatures typical of the 2-cell, 8-cell, ICM/epiblast, TE and PrE lineages (Supplementary Table 1 and Supplementary Figure 3c) to classify all cells in this dataset (Figure 3c,d). This analysis suggested that the E3.5 miR-203-deficient cell cluster was composed of 8-cell-like and ICM/epiblast cells, as well as a small number of 2-cell-like cells that were not present in other groups. Pseudotime inference in this group of cells confirmed the expected transition from 2C-to 8C-like cells before giving rise to the three major embryonic lineages (Figure 3e-g and Supplementary Figure 3d,e). By E4.5, all control, *miR-203*(+/tet)^Dox^ and miR-203-deficient embryos had separated populations representing TE, as well as the two lineages generated in the second cell fate decision in the embryo: the epiblast and PrE. However, the ratio in the number of cells belonging to these populations was altered, with miR-203-deficient and *miR-203*(+/tet)^Dox^ embryos having a lower proportion of TE cells (Figure 3h). Interestingly, *miR-203*(*−*/*−*) E3.5 embryos showed a significant overexpression of MERVL transcripts when compared to wild-type or *miR-203*(+/tet)^Dox^ embryos, and these differences were still evident in E4.5 embryos (Figure 3i and Supplementary Figure 3f). MERVL high-cells were mostly found in the 2-cell- and 8-cell-like clusters described above in the *miR-203*(*−*/*−*) E3.5 embryos (Figure 3j and Supplementary 3g,h), but were also present in some *miR-203*(*−*/*−*) and *miR-203*(+/tet)^Dox^ Epiblast-like clusters at E4.5 (Figure 3j). In agreement with the increased levels of 2-cell-like markers, *miR-203*(*−*/*−*) embryos displayed decreased expression of pluripotency markers by E3.5 (*Nanog, Oct4, Sox2;* Supplementary Figure 3i). However, by E4.5 pluripotent transcripts recovered and were more abundant in the epiblast lineage of miR-203-null embryos. These mutant cells displayed decreased abundance of TE markers and higher expression of hypoblast markers, in agreement with the differences in the representation of those cells in mutant embryos (Figure 1h).

### miR-203 modulates the P300 histone acetyltransferase during early developmental stages

The analysis of pathways differentially expressed in E3.5 embryos identified chromatic modifying enzymes as the most downregulated pathway in miR-203 overexpressing embryos (Figure 4a) suggesting a possible effect of miR-203 in the transcripts encoding these enzymes. Downregulated transcripts encoding chromatin regulators in *miR-203*(tet)^ON^ embryos included *Ep300, Pbrm1, and Smarcd1*, among others (*Atf7ip, Kmt2a, Kat6a, and Tada2b*; Figure 4b). On the other hand, these three transcripts were upregulated in *miR-203*(*−*/*−*) embryos, together with *Kat6b, Arid1a, Kdm6a and Dr1*; Figure 4c). We tested the ability of miR-203 in directly repressing these transcripts by generating reporter constructs with the 3’-UTR of each of them downstream of a luciferase reporter. Expression of miR-203 was able to repress most of these transcripts (Figure 4d) with the possible exception of *Arid2* and *Atf7ip*, two chromatin regulators whose transcript levels were also reduced although did not reach statistical significance in the differential expression analysis. Mutation of the predicted miR-203 binding sites of these luciferase-3’-UTR reporters prevented miR-203-driven repression (Figure 4d), suggesting a direct control of these transcripts by miR-203.

**Figure 4.**
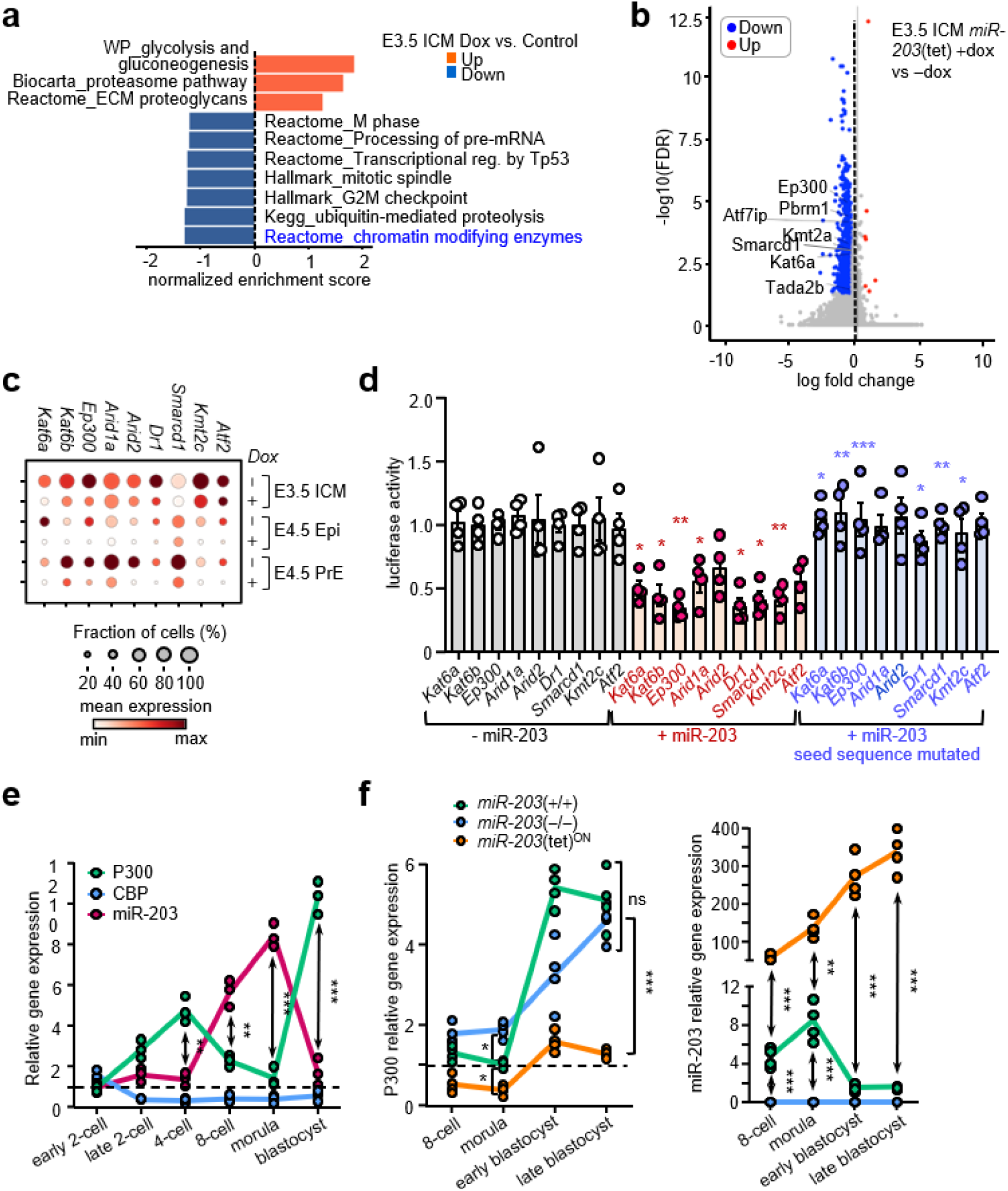
Histone acetyltransferase enzymes are direct biochemical targets of miR-203. **a**, Top categories in the Gene Ontology analysis of the genes significantly up- and downregulated in the ICM of E3.5 blastocysts developed in utero in the presence or absence of doxycycline to conditionally induce *miR-203* expression. Note that the go-terms for chromatin modifying enzymes are amongst the most downregulated in these samples. **b**, Volcano plot displaying the gene expression changes in E3.5 blastocysts overexpressing miR-203 compared to wild-type controls. Note that several genes encoding members of the histone acetyltransferase enzymes family are downregulated, especially *Ep300*. **c**, Dot plot depicting the expression of individual genes contained in the Go-terms for histone modifying enzymes in embryos from the stated genotypes and treatments (*+* dox: *+; −* dox: *-).* Scale bar represents the mean gene expression. **d**, Bar plot displaying luciferase activity (fold increase in arbitrary units) in HEK293T cells transfected with the corresponding luciferase vectors containing a wildtype or mutated seed sequence within the 3’UTR region of the predicted miR-203 target transcripts. Each data point represents a biological replicate of the assay, and error bars indicate SEM. Magenta asterisks represent the comparisons between the – miR-203 and + miR-203 groups; and blue asterisks the comparisons between the + miR-203 and + miR-203/seed sequence mutated groups. **e**, Gene expression graph showing the expression levels of the genes *Cbp* and *Ep300* (respect *Hprt* as a housekeeping control) and miR-203 (respect miR-16 as housekeeping control miRNA). Note the antagonistic pattern of gene expression displayed between *Ep300* and *miR-203,* and that the expression of *Cbp* remains stably low during preimplantation development. **f**, Relative expression of *Ep300* respect *Hprt* as a housekeeping control in early embryos of the indicated genotypes. Note the different levels of *Ep300* transcripts upon genetic manipulation of *miR-203* within the morula stage and the clearance of these differences with the course of development. **g**, Relative expression of *miR-203* within embryos isolated in the different preimplantation stages stated. In **e**, **f**, **g** each data point represents one biological replicate containing a minimum of 20 embryos of the indicated stage. Data in **d**-**g** are analyzed by a 1-way ANOVA with Tukey multiple comparisons post-hot test. *, *P*<0.05, **, *P*<0.01, ***, *P*<0.001, ****, *P*<0.0001, ns: non-significant.

Among the validated candidates, we observed that P300, but not the related acetyltransferase CREB binding protein (CBP), displayed an expression pattern inverse to that of miR-203, with increasing levels during the 2-4-cell stages and low levels at the 8-cell and morula stages when miR-203 peaks (Figure 4e), in agreement with previous studies (Wang *et al*, 2022). Interestingly, expression of P300 increased after miR-203 was silenced in normal blastocysts, and upregulation did not happen when miR-203 expression was maintained high in the miR-203(tet)^ON^ model (Figure 4f).

### miR-203 regulates developmental timing by controlling acetyltransferase levels in preimplantation embryos

To test the relevance of P300 as a miR-target we made use of Trichostatin A (TSA), a molecule that induces the acetylation and stability of P300, in E2.5 and E3.5 embryos isolated from the eE-Reporter mouse. As depicted in Figure 5a and Supplementary Figure 4a, the P300 stabilizer TSA induced a phenotype similar to that observed in *miR-203*(*−*/*−*) embryos, including developmental delay and strong expression of MERVL at later developmental stages. The effect of TSA was reproduced with suberoylanilide hydroxamic acid (SAHA), a pan-histone deacetylase (HDAC) inhibitor (Supplementary Figure 4b). On the other hand, A485, a catalytic inhibitor of the P300 and CREB binding protein (CBP) acetyltransferases, also induced a delay in embryo development (Figure 5a and Supplementary Figure 4a). Importantly, this P300/CBP inhibitor prevented the expression of MERVL in *miR-203*(*−*/*−*) embryos (Figure 5a and Supplementary Figure 4a), suggesting that these acetyltransferases were required for MERVL expression in the absence of miR-203. Similarly, short interfering RNAs (siRNAs) against *Ep300* also displayed a significant effect in delaying preimplantation embryonic development, and this effect was reproduced to a lesser extent after knock down of the lysine acetyltransferases *Kat6a* or *Kat6b* (Figure 5b).

**Figure 5.**
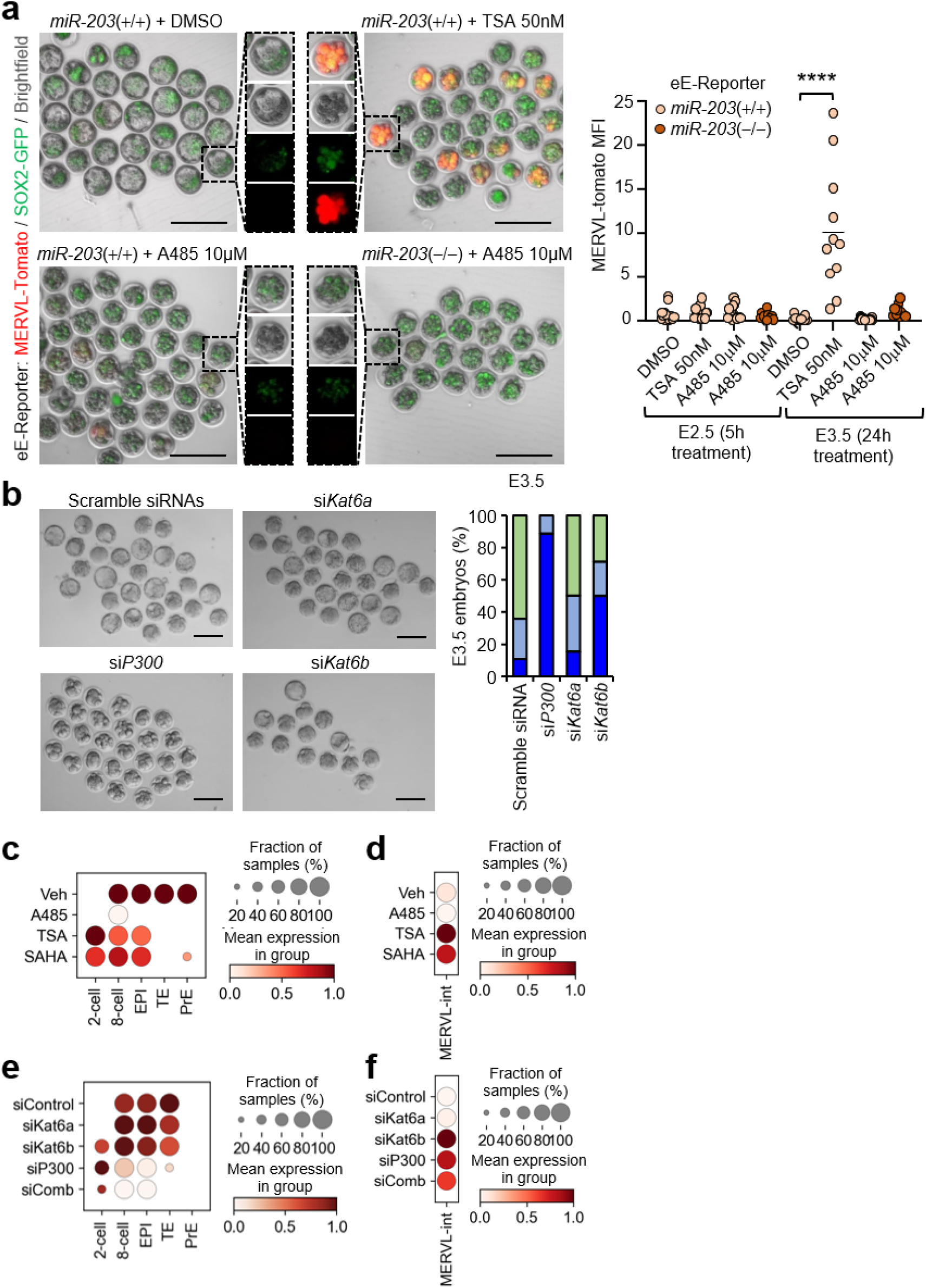
Direct manipulation of H3K27 histone acetyltransferase and deacetylase activity mimics miR-203 phenotypes in early embryos. **a**, Brightfield and fluorescence micrographs of live embryos of the stated genotypes isolated at E2.75 and treated with the indicated small molecule inhibitors for 24h. Plot to the right depicts the quantification of the MERVL-tomato mean fluorescence intensity per live embryo (MFI) after 5 and 24h of treatment. **b**, Brightfield micrographs of preimplantation embryos injected with the indicated siRNAs at the zygote stage (E0.5) and developed in vitro for 3 days. Scale bar: 100µm. The bar plot to the right shows the quantification of the percentage of embryos in each of the embryonic stages listed after microinjection with siRNAs at the zygote stage (E0.5) and their development in vitro for 72h. **c-f**, Dot plots depicting the expression of the stated gene signatures (c, e) or MERVL sequences (d, f) for the indicated experimental conditions. Scale circles and bars represent the fraction of samples and mean gene expression per group, respectively. Data in **a** are analyzed by a 1-way ANOVA with Tukey multiple comparisons post-hot test. ****, *P*<0.0001.

To analyze more in detail the molecular effects of these treatments, we exposed E2.5 8-cell stage embryos with the above-mentioned small molecule inhibitors for 24h in vitro, and samples were processed for low input RNA-seq. Stabilization of P300/CBP or HDAC inhibition resulted in significant increase in cells displaying a 2-cell-like transcriptional program (Figure 5c and Supplementary Figure 4c), including the expression of MERVL transcripts (Figure 5d). The effect of specifically modulating P300 levels was validated by microinjecting zygotes with siRNAs against P300, 72 h before harvesting samples for RNAseq analysis. Knock down of *Ep300* resulted in over-representation of 2-cell-like transcripts (Figure 5e and Supplementary Figure 4d) as well as overexpression of MERVL sequences (Figure 5f). Interestingly, these effects we are also reproduced after known down of *Kat6b*, suggesting that miR-203 may modulate preimplantation developmental timing by regulating P300 and KAT6B levels in preimplantation embryos.

We finally tested the effect of miR-203 levels in lysine acetylation at H3K9 and H3K27, two sites involved in the activation of transcripts, by testing the genome-wide profile of acetylation of these sites using low input bulk CUT&RUN on E2.5 disaggregated embryos (Meers *et al*, 2019; Wang *et al*., 2022). Lack of miR-203 resulted in increased acetylation of genes belonging to the 2-cell-like signature whereas overexpression of miR-203 had a small effect on reducing acetylation of these gene promoters (Figure 6a). Statistical analysis using gene set enrichment indicated increased H3K9ac and H3K27ac marks in 2-cell genes as well as MERVL sequences in miR-203-null embryos (Figure 6b,c). All these data suggest that lack of miR-203 induces increased levels of acetyltransferases such as EP300 and possibly KAT6B, thereby resulting in increased acetylation and expression of 2-cell stage transcripts during the 8-cell-to blastocyst stages of preimplantation development.

**Figure 6.**
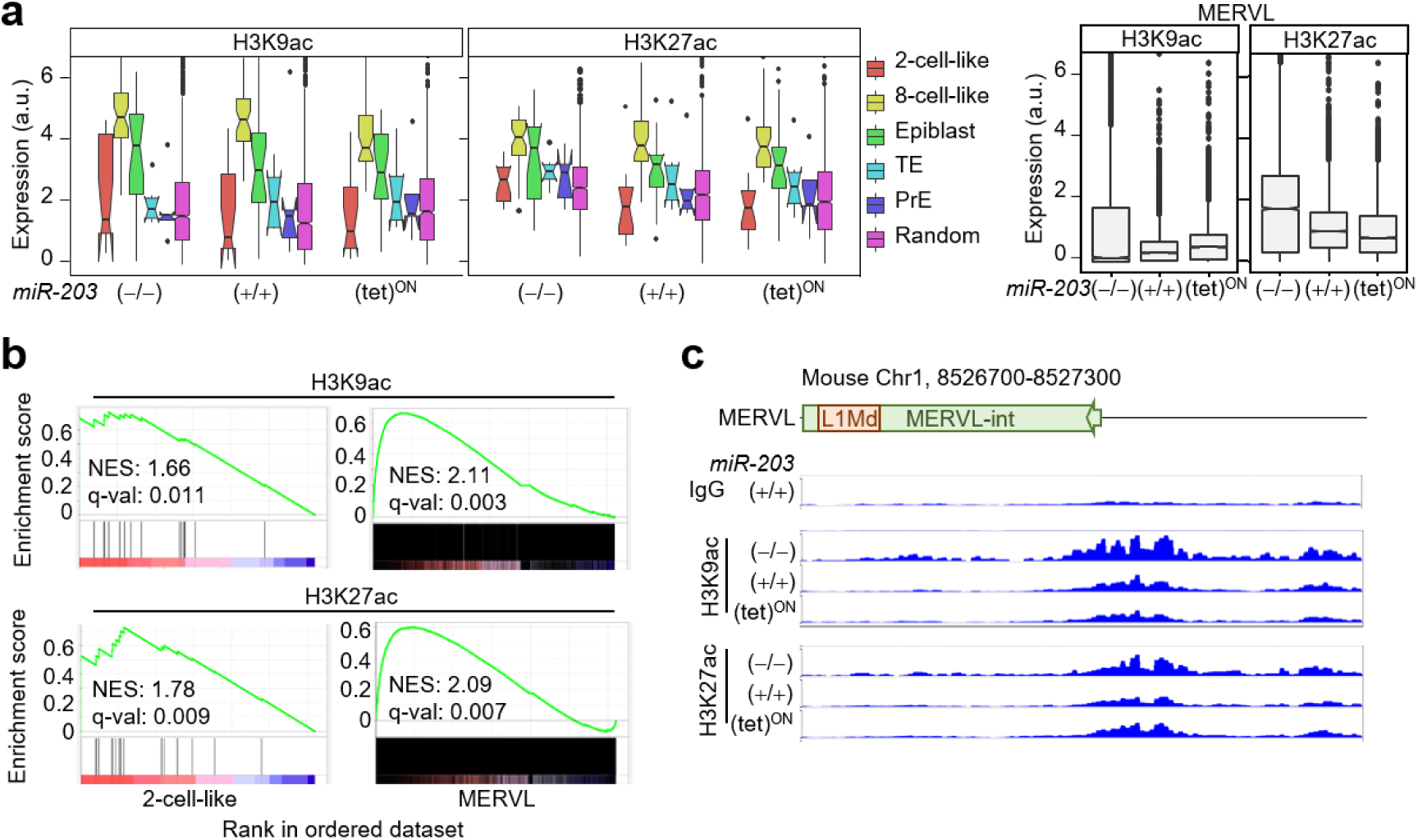
Loss of miR-203 promotes increased histone acetylation in totipotency-associated genetic loci. **a**, Box plot depicting the levels of histone H3 lysine 9 acetylation (H3K9ac) and histone H3 lysine 27 acetylation (H3K27ac) in genomic loci corresponding to genes contained in the signatures listed to the right within E2.75 embryos of the indicated genotypes and treatments. **b**, Gene enrichment profiles corresponding to H3K9ac and H3K27ac histone marks in 2-cell-like and MERVL gene signatures from the embryos analyzed in a. **c**, Genome view analysis of the indicated histone acetylation marks within the chromosomal region corresponding to one specific MERVL sequence. Note the increase in H3K9 and H3K27 acetylation upon miR-203 loss.

## Discussion

From early embryogenesis to the assembly of neural circuits, development relies on precise timing (Ebisuya & Briscoe, 2018; Rayon, 2023). During embryogenesis, a series of events ought to take place in the correct order and at a given speed of progression in a developmental sequence to ensure a correctly paced development. In early mammalian embryogenesis, preimplantation development entails the coordination of several cellular processes including cell polarization, cell compaction and cell fate restriction concomitant with lineage segregation (Menchero *et al*., 2017; Wilkinson *et al*., 2023; Zhu *et al*, 2020).

Epigenetic programs are known to control developmental tempo in a variety of biological scenarios including neural differentiation (Ciceri *et al*., 2024) and preimplantation embryogenesis (Creyghton *et al*, 2010; Dahl *et al*, 2016; Wang *et al*., 2022). Several epigenetic regulators including the histone acetyltransferase and transcriptional coactivator P300 have been pinpointed as critical for these processes (Chan *et al*, 2019; Ferrie *et al*, 2024). Most of the upstream regulators behind these epigenetic programs have remained elusive, and our findings help account for the role of the micro-RNA miR-203 in regulating the expression of totipotency-associated genes via the activity of MERVL endogenous retroviral repeats. MERVL transcripts are among the first transcripts expressed during the early zygotic activation at the 2-cell stage, as their UTR sequences act as functional promoters of several known totipotency genes (Macfarlan *et al*., 2012). In line with these findings, its activation in pluripotent cells is sufficient to induce a transient 2C-like cell state (De Iaco *et al*, 2017; Hendrickson *et al*, 2017; Yang *et al*, 2020). MERVL sequences function as chromatin organizers in early embryos (Asimi *et al*, 2022) and full-length MERVL transcripts, but not encoded retroviral proteins, are essential for preimplantation development (Sakashita *et al*, 2023).

The epigenetic control of cell fate transitions during preimplantation development in mammals is well established, although the exact mechanisms controlling changes in time and space remain unclear. Several mechanistic studies suggested some microRNAs to be involved in the epigenetic control of MERVL expression in in vitro cultures of embryonic stem cells, including miR344 and miR-34a (Choi *et al*, 2017; Yang *et al*., 2020). These multiple mechanisms of control also suggest high redundancy. It is well established that microRNAs fine-tune gene expression, rather than being the primary determinant of expression levels. By using an integrative and combinatorial approach, multiple miRNAs may target an individual gene and an individual miRNA may target multiple genes, making it very difficult for almost any biological process to escape from miRNA regulation (DeVeale *et al*, 2021). For instance, no differences in the levels of MERVL and 2C transcripts were observed in miR-34a-deficient preimplantation embryos, suggesting highly redundant molecular networks mediating MERVL silencing (Choi *et al*., 2017). In addition to the presence of multiple miR-34 family members, it is also conceivable that multiple microRNAs could act redundantly to repress MERVL expression to ensure rapid and efficient MERVL silencing in vivo. Genetic depletion of miR-203 in early embryos however yields MERVL hyperactivation phenotypes and a global protraction of developmental timing, likely due to the fact that miR-203 is an orphan family member whose genetic loss is less easily compensated. Whereas microRNA activity is dispensable for pre-implantation development (Bernstein *et al*, 2003; Wang *et al*, 2007), homozygous mutations in genes encoding many epigenetic regulators cause early developmental failure (Grosswendt *et al*, 2020), suggesting additional layers of regulation beyond these non-coding RNAs.

The fact that heterozygous or hemizygous mutations in the genes encoding the acetyltransferases P300 and CBP cause the Rubinstein-Taybi syndrome, characterized by distinctive facial features and varying degrees of intellectual disability (Roelfsema *et al*, 2005; Van Gils *et al*, 2021), suggests that P300 is happloinsufficient for cell fate decisions in development. In fact, even a relatively small imbalance in these proteins has significant developmental effects (Roelfsema *et al*., 2005). P300 and CBP act as integrators of the signals from various molecular pathways that compete for these factors to determine chromatin accessibility (Goodman & Smolik, 2000). For instance, P300 is known to participate as a SOX2 co-activator in pluripotent cells (Yoo *et al*, 2023), as well as a mediator of trophoblast stem cell pluripotency (Dou *et al*, 2024).Our data suggest that miR-203 may participate in regulating P300 levels during preimplantation development. In the absence of miR-203, P300 levels would rise, allowing its binding to additional genomic loci such as those involved in the 2C stage (including MERVL), allowing the expression of both totipotency and pluripotency genes, and increasing lineages other than TE, as observed in *miR-203*(*−*/*−*) embryos. On the other hand, overexpression of miR-203 would lead to a general deficiency in the availability of P300 for totipotency and pluripotency enhancers, leading to specific defects in later lineage determination. Whereas these hypotheses remain speculative, our data along with previous observations reporting the control of DNMT3A/B by miR-203 in pluripotent cells (Salazar-Roa *et al*., 2020), suggest new functions for this microRNA in early developmental transitions. Given the increasing relevance of assisted reproductive technologies, these data may have implications for controlling the regenerative potential of pluripotent cells for medical needs (Gonzalez-Martinez & Malumbres, 2022).

## Methods

### Mouse models

The miR-203 conditional knockout model was generated by flanking the *mmu-mir203* locus with loxP sites (Genebridges) using homologous recombination into ES cells (Supplementary Figure 1b). A frt-flanked neomycin-resistant cassette was used for selection of recombinant clones. Elimination of the neomycin-resistant cassette was achieved using a CAG-Flpe transgenic line (Rodriguez *et al*, 2000) and loxP sites were recombined using a EIIa-Cre transgene (Schwenk *et al*, 1995) resulting in the *miR-203*(*−*/*−*) allele (Figure 1b and Suppl. Figure 1b). The miR-203 inducible model (Salazar-Roa *et al*., 2020) and Sox2-GFP transgenic allele (Suh *et al*., 2007) were described previously. For the generation of the MERVL-Tomato transgenic allele, the expression vector reported previously (Macfarlan *et al*., 2012) was isolated from a bacterial extract using the EndoFree Plasmid Maxi Kit (Qiagen #12362), digested with SacII for linearization, phenol/chloroform treated and resuspended in microinjection buffer 10 mM Tris-HCl pH 7.5, 0.1 mM EDTA at 1 ng/*μ*l. Transgenic mice were produced by pronuclear injection of the linearized vector into zygotes obtained from superovulated B6.CBA females crossed with males containing the *Col1a1_tetO-miR203* and *Rosa26*_*rtTA* alleles (Salazar-Roa *et al*., 2020) using standard protocols. Out of 66 mice born, 10 were confirmed positive for transgene integration. Founders were crossed with C57BL6/J females to test for germ line transmission.

Mouse studies were performed using a B6.CBA genetic background under specific-pathogen-free conditions at the Spanish National Cancer Research Center (CNIO) animal facility, following the animal care standards of the institution. The animals were observed daily, and diseased mice were humanely euthanized in accordance with the Guidelines for Humane Endpoints for Animals Used in Biomedical Research (Directive 2010/63/EU of the European Parliament and Council and the Recommendation 2007/526/CE of the European Commission). All animal protocols were approved by the committee for animal care and research of the Instituto de Salud Carlos III-Comunidad de Madrid (Madrid, Spain).

### Mice superovulation, embryos isolation, and microinjection

For female mice superovulation, 4 to 5 weeks old females were superovulated by intraperitoneal injection of 5 IU of pregnant mare serum gonadotropin per female (PMSG; Sigma G4877-1000IU or ProSpec #HOR-272) on Day 1 at 1:00 p.m. On day 3 at 12:00 p.m, females were injected intraperitoneally 5IU of human chorionic gonadotropin (hCG; sigma C1063-10VL) and mated with the desired males (8 to 20 weeks old). The next day vaginal plugs were assessed at 7-8 am (0.5dpc or E0.5). Zygotes were collected at E0.5, 2-cell stage embryos at E1.5, 8-cell stage embryos at E2.5, morulae at E2.75, and blastocysts at E3.5 (mid blastocyst stage) or E4.5 (late blastocyst stage). Isolation of zygotes was achieved by dissecting out the cumulus zygotes complexes and briefly treating them with type IV hyaluronidase (Merck #H4272), whereas 2-cell, 8-cell and morula stage embryos were obtained by flushing the oviduct with M2 medium (Sigma-Aldrich # M7167) by inserting a fine needle through the infundibulum and collecting the embryos. Blastocysts were obtained by flushing the uterus at E3.5/4.5. After isolation, embryos were processed for analysis or cultured in 20uL drops of KSOM EmbryoMax medium (Sigma-Aldrich #MR-106-D) carefully covered by mineral oil suitable for embryo culture (lifeguard oil #LGUA-5 00) in P35 culture dishes. For embryo microinjection, E0.5 zygotes or E1.5 2-cell stage embryos were isolated, and siRNAs or DNA constructs diluted in microinjection buffer (10 mM Tris pH 7.4, 0.1 mM EDTA) in the indicated concentrations were microinjected in the cytoplasm or the pronucleus, respectively. When needed, the resulted microinjected embryos were transferred to Crl:CD1(ICR) pseudo-pregnant females.

### Analysis of microRNA expression levels in preimplantation embryos

For real time PCR, embryos were harvested and pipetted into 8uL of cell lysis buffer from the TaqMan MicroRNA Cells-to-CT Kit (Invitrogen #4391848) in a 200 μL low binding tube, pipetted up and down 10 times with low binding pipette tips, and incubated for 8min at room temperature. An average of 15-20 embryos per experimental condition and replicate were used. After incubation, 1 μL of stop buffer was added to the mix, which was pipetted up and down 5 times and kept on ice. After miRNA isolation, miRNA samples were shortly subjected to cDNA synthesis and real time PCR amplification with two-step miRNA assays from TaqMan probes following manufacturer instructions, using the TaqMan Universal PCR Master Mix, no AmpErase UNG (ThermoFisher #4364343) in a QuantStudio 5 real-time PCR system. The Taqman two-step miRNA expression assays used in this study were the miR-203 pri-miRNA assay (Assay ID: Mm03306530_pri); the miR-16 pri-miRNA assay (Assay ID: Mm03306798_pri); the miR-203 miRNA assay (Assay ID: 000507), and the miR-16 miRNA assay (Assay ID: 000391).

### Single-cell RNA sequencing

For single-cell RNA sequencing of E3.5 blastocysts, zona pellucida was removed passing the embryos by mouth pipetting through three consecutive drops of Tyrode’s acidic solution pH 2.5 (Sigma T1788). After careful examination under the microscope for disintegration of the pellucida, tyrode’s solution was immediately neutralized with a double volume of PBS pH 7.4 (Gibco #10010023), and embryos were placed into 20 μL of dissociation buffer composed of TrypLE™ Express Enzyme (Gibco # 12604013) / Accutase (Gibco # A1110501) in a 1:1 ratio, in a low binding 200 μL microcentrifuge tube. To avoid attachment of the embryos to the pulled crystal capillaries, capillaries were coated with 1% BSA in PBS beforehand. Blastocysts were incubated for 20 minutes in dissociation buffer at 37°C, pipetted up and down gently 60 times with 20 μL low binding pipette tips, incubated 10 additional minutes at 37°C and pipetted up and down again gently 40 times to disaggregate the last ICM cell clumps. After dissociation, the resulting cell suspension was neutralized with 40uL of 1% cell culture grade BSA (Sigma # A8806). In the case of 8-cell and morula stage embryos, pellucida was removed as abovementioned and embryos were placed in 20 μL of Accutase (Gibco # A1110501) for 20 min at 37°C. After incubation, embryos were pipetted up and down 50 times until complete disaggregation and neutralized by the addition of 40 μL of 1% cell culture grade BSA (Sigma # A8806) to avoid the formation of clumps after dissociation. Then, cell suspensions were immediately placed on ice until processing for scRNA-seq. For scRNA-seq analysis, cell samples were loaded onto a 10x Chromium Single Cell controller chip B (10x Genomics) following manufacturer’s intructions (Chromium Single Cell 3’GEM, Library & Gel Bead Kit v3, ref. PN-1000075). Intended targeted cell recovery of ∼1000 - 10000 cells depending on the embryonic stage. Generation of gel beads in emulsion (GEMs), barcoding, GEM-RT clean-up, cDNA amplification and library construction were all performed as recommended by the manufacturer. scRNA-seq libraries were sequenced with an Illumina NextSeq 550 (using v2.5 reagent kits). Raw images generated by the sequencer are submitted to analysis, per-cycle base calling and quality score assignment with Illumina’s Real Time Analysis integrated primary analysis software (RTA v2). Conversion of BCL (base calls) binary files to FASTQ format is subsequently performed with bcl2fastq2 (Illumina).

### Ultra-low input bulk RNA-seq

For ultra-low input RNA-seq, 15-20 embryos were collected per sample and resuspended in 8 mL of low input lysis buffer of the NEBNext Single Cell/Low input cDNA Synthesis & Amplification module (#E6421S). Whole cells were processed with the “NEBNext Single Cell/Low Input RNA Library Prep” kit (NEB #E6420) by following manufacturer instructions. Double stranded cDNA production was performed by limited-cycle PCR. Sequencing libraries were completed with the “NEBNext Ultra II FS DNA Library Prep Kit for Illumina” (NEB #E7805) and subsequently analysed on an Illumina instrument by following manufacturer’s protocols. The resulting purified cDNA libraries were applied to an Illumina flow cell for cluster generation and sequenced on Illumina NovaSeq X (with NovaSeq X Series Reagent Kits) by following manufacturer’s protocols using PairEnd (2×151 bases) in KO experiment and Illumina NextSeq 550 (with 75 cycle High Output v2.5 reagent kit) using SingleRead (86 bases) in pharmacological modulation experiments. Raw images generated by the sequencer are submitted to analysis, per-cycle basecalling and quality score assignment with Illumina’s Real Time Analysis integrated primary analysis software (RTA4). Conversion of BCL (base calls) binary files to FASTQ format is subsequently performed with BCL Convert (Illumina).

### Computational analysis of transcriptomic data

For ultra-low input RNAseq, we used the cluster_rnaseq pipeline (https://github.com/cnio-bu/cluster_rnaseq/tree/master). Sequencing quality control was performed using multiqc preprocessing (v1.18; https://multiqc.info), and adapters were removed with BBDuk (v39.01, https://github.com/BioInfoTools/BBMap/blob/master/sh/bbduk.sh). For each sample, we performed read alignment to the mm10 version of GRCh38 mouse reference genome using STAR (v2.7.11a) and quantification using featureCounts from Subread package (v2.0.6; (Liao *et al*, 2019)) and count matrices were used for downstream analysis in scanpy v. 1.8.2. We used TEtranscripts (v2.2.3; https://github.com/mhammell-laboratory/TEtranscripts) in order to quantify TE elements in our low input RNA samples using bam files extracted with STAR in aligment step, annotation file for mm10 version of GRCh38 mouse reference genome and for transcript elements of GRCh38 mouse reference pre-generated by RepeatMasker 3.0 (http://www.repeatmasker.org).

For single-cell RNAseq data, quality control was performed using fastqc (version 0.11.9). For each sequenced single-cell library, we performed read alignment to the mm10 version of GRCh38 mouse reference genome. Quantification and initial quality control were performed using STARsolo of STAR (version 2.7.10a) and STARsolo filtered count matrices were used for downstream analysis. We integrated the filtered count matrices from STARsolo and analysed them with scanpy v. 1.8.2 following their recommended standard practices (Wolf *et al*, 2018). In brief, we excluded genes expressed by less than three cells, excluded cells expressing fewer than 150 genes, and cells with more than 15% mitochondrial and 25% ribosomal content. Expression data was normalized and converted to log(X + 1). Next, for dimensionality reduction we identified highly variable genes, scaled the data and calculated PCA to observe the variance ratio plot and decide on an elbow point which defined n_pcs. This was used for calculating neighbourhood graph, UMAP and further Louvain clustering. RNA velocity was used to investigate directed dynamic information by leveraging splicing kinetics using scVelo v. 0.2.4 (La Manno *et al*, 2018). For the identification of transposable elements we used scTE v. 1.0 (He *et al*, 2021) following the same parameters used with STARsolo.

The origin of gene signatures used for signature scores in shown in Supplementary Table 1. All differential gene expression analysis were performed with scanpy using t-test method for comparisons between subpopulations and conditions. Only significant differentially expressed genes were considered for downstream analysis (FDR (bonferroni) < 0.05). GSEA Preranked was used to perform gene set enrichment analysis for the selected gene signatures on a pre-ranked gene list, setting 1,000 gene set permutations (Subramanian *et al*, 2005) using the Python wrap (Fang *et al*, 2023). Only those gene sets with significant enrichment levels (FDR < 0.25) were considered.

### Luciferase assays

Luciferase assays were performed in HEK293T cells. Briefly, 2.10^5^ cells per well were seeded on 6-well plates, and in 24h cells were transfected using Lipofectamine 2000 (Invitrogen), following the manufacturer’s instructions. The 3′-UTR regions from the murine genes *Kat6a, Kat6b, Ep300, Arid1a, Arid2, Dr1, Smarcd2, Kmt2c* and *Atf7ip* were amplified by PCR with specific primers (Supplementary Table 2) using mouse cDNA from ES cell cultures. PCR products were verified by sequencing cloned into the pGL3-Control vector (Promega) by Gibson Assembly, downstream of the luciferase reporter gene. Mutations in the miR-203 seed sequence binding sites were generated by site-directed mutagenesis (Supplementary Table 3) and subsequently verified by sequencing. Overexpression of miR-203 or control GFP was achieved by transfection with either pMCSV-GFP or pMCSV-miR-203-GFP vectors (Salazar-Roa *et al*., 2020), in combination with the pGL3-derived vectors, and *Renilla* as a control. Luciferase measurement was achieved 48h post-transfection using a luminescence microplate reader (Bio-Tek).

### Ultra-low input CUT&RUN

CUT&RUN samples were generated following the original (Meers *et al*., 2019) and modified protocols (Wang *et al*., 2022) for early embryos. For CUT&RUN, 100 disaggregated blastocysts were used per sample. For each sample, 10 μl of BioMag Plus Concanavalin A beads (Polysciences # 86057-3) were washed twice in 500 μL of Binding Buffer (20 mM HEPES-KOH pH 7.9, 10 mM KCl, 1 mM CaCl_2_, 1 mM MnCl_2_), and after buffer removal were resuspended in 10 μl Binding Buffer. Then, 8-cell stage embryos or disaggregated blastocysts (as stated in the scRNA-seq section) were added to 45μl Wash Buffer (150 mM NaCl, 0.5 mM Spermidine, 20 mM HEPES-NaOH pH 7.5, 1X Roche Protease Inhibitor Cocktail), mixed with the 10μl Binding Buffer containing magnetic beads, gently tapped twice, and incubated for 10 min at room temperature in a 200 μL low binding tube. Buffer was removed by placing the tube on a magnet stand, the beads were resuspended in 50 μl antibody Incubation Buffer (1X Wash Buffer with 0.02% Digitonin and 2 mM EDTA) with the indicated antibody and kept at 4°C overnight. Then, samples were washed twice with 100 μl Wash Buffer with 0.02% Digitonin. The beads were resuspended in 100 μl Wash Buffer with 0.02% Digitonin and 500 ng/ml CNIO home-made pA-MNase, and rotated for 2h at 4°C. Beads were then washed twice with 100 μl of Wash Buffer with 0.02% Digitonin, resuspended in 100 μl ice-cold Wash Buffer with 0.02% Digitonin and 2 μl 100 mM CaCl_2_, and incubated in ice in a metallic rack for 30 min. The reaction was stopped by adding 12 μl of 10X Stop Buffer (1.700 mM NaCl, 100 mM EDTA, 20 mM EGTA, 0.02% Digitonin, 250 μg/ml RNaseA, 250 μg/ml Glycogen). The tube was then incubated at 37°C for 15 min to release the CUT&RUN fragments into the supernatant. After spinning the sample for 1min and putting it in a magnetic stand, the supernatant was transferred to a new 1.5 ml ml low binding tube while discarding the beads. DNA was purified by fist adding 2.5 μl 10% SDS, and 2.5μl Proteinase K (20 mg/ml) to the supernatant and incubating it at 55°C for 1 h. GlycoBlue (Invitrogen #AM9516) was added to the supernatant before the supernatant was mixed with 200 μl Phenol:Chloroform:Isoamyl Alcohol (Invitrogen) by vortexing. Samples were then centrifuged at 16000 g for 5min at room temperature. After carefully transferring the supernatant to a new 1.5 ml LoBind tube with low binding pipette tips, 26 μl 3M Sodium Acetate, 1 μl Glycogen (20 mg/ml), and 750 μl cold 100% Ethanol were added, mixed by vortexing, and incubated at -20°C for 1h. Then, the tube was centrifuged at 16,000 g for 20 min at 4°C and the supernatant was discarded. The precipitated DNA pellet was washed with 500uL room temperature 70% Ethanol, and then dried at 55°C for 15min. Finally, the DNA pellet was dissolved overnight in 20 μl ultrapure H_2_O. DNA samples were processed through subsequent enzymatic treatments for end-repair, dA-tailing, and ligation to adapters with “NEBNext Ultra II DNA Library Prep Kit for Illumina” (NEB #E7645). Adapter-ligated libraries were completed by a PCR of 21 cycles of amplification, with 63° T(ann/ext) for 10s, and extracted with a (single) double-sided SPRI size selection. Libraries were applied to an Illumina flow cell for cluster generation and sequenced on an Illumina NextSeq550 in paired-end 43-bp mode by following manufacturer’s recommendations.

Quality control was performed using fastqc. Paired-end reads were aligned against mm10 genome using Bowtie2 (v.2.4.2) with the following parameters “-local --very-sensitive-local --no-unal --no-mixed --no-discordant --phred33 -I 10 -X 700”. Then we excluded reads that fail in the alignment using samtools (v.1.11) and duplicated reads with GATK (v.4.1.9). Finally, the bam files were converted to bigwig using bamCoverage from deeptools (v.3.5.1). To extract the signal of the promoters, we quantified the amount of signal in a 2.5kb window centered in the TSS of the genes and normalized the values using the median value of the 5000 more expressed genes for each condition (Dahl *et al*., 2016). Additionally, we quantified the signal in the MERVL sequences and normalized them using the value previously calculated. GSEA preranked (v.4.3.2) was used to evaluate the enrichment in the different conditions compared to controls.

## Statistics

Statistics was performed using Prism 9 software (GraphPad Software) or Python 3.8. All statistical tests of comparative data were done using an unpaired, 2-tailed Student’s t test with Welch’s correction, or 1-way ANOVA with Tukey’s multiple-comparison test when appropriate. Data are expressed as the mean of at least 3 independent experiments ± SEM; with a *P* value of less than 0.05 considered statistically significant.

## Data availability

Data from transcriptomics and CUT&RUN assays will be accessible from Gene Expression Omnibus (GEO). Python notebooks used in this analysis are available at https://github.com/malumbreslab/miR203_developmental_timing.

## Acknowledgements

The authors wish to thank Maria J. Bueno (CNIO) and Marta Gómez de Cedrón (IMDEA Madrid) for the generation of the miR-203-null allele, the Genomics and Mouse Genome Editing Units at CNIO for their outstanding technical work and scientific support, and Miriam García for an excellent mouse colony management at the CNIO animal facility. A.S.B. was supported by a FPI contract from the Spanish Ministry of Science and Innovation (MICINN; PRE2022-103179). A.G. received a predoctoral contract from the ”la Caixa” Foundation (LCF/BQ/DI21/11860054). M.M. lab was supported by research grants from MICINN/AEI/FEDER (PID2021-128726 and PDC2022-133408-I00), and Comunidad de Madrid (Y2020/BIO-6519 and S2022/BMD-7437). VHIO would like to acknowledge the Cellex Foundation for providing research facilities and equipment and the CERCA Program from the Generalitat de Catalunya for their support on this research. CNIO (CEX2019-000891-S) and VHIO (CEX2020-001024-S/AEI/10.13039/501100011033) are Centers of Excellence Severo Ochoa (Agencia Estatal de Investigación, MICINN).

## Author contributions

J.G.M.: Conceptualization, Methodology, Formal analysis, Investigation, Supervision, Writing; A.S.B.: Methodology, Formal analysis, Investigation, Writing – Review & Editing; E.A.: Methodology, Formal analysis, Investigation; A.G.: Methodology, Formal analysis, Investigation; D.G.: Methodology, Formal analysis, Investigation; J.M.; Methodology, Investigation; A.L.: Methodology, Investigation; S.O.: Methodology, Formal analysis, Investigation, Writing – Review & Editing; M.M.: Conceptualization, Methodology, Formal analysis, Resources, Supervision, Writing, Project administration and Funding acquisition.

