## Supplementary Information for "miR-203 controls developmental timing and early fate restriction during preimplantation embryogenesis"

by José González-Martínez et al.

### Table of Supplementary Material

#### Supplementary Figures

**Supplementary Figure 1.** miR-203 expression levels in vivo and generation of miR-203 mutant alleles.

**Supplementary Figure 2.** The eE-Reporter mouse registers the activity of MERVL endogenous retroviral elements associated with the totipotent state during early mouse embryogenesis.

**Supplementary Figure 3.** Analysis of cell lineages in miR-203-mutant embryos.

**Supplementary Figure 4.** Pharmacological modulation histone acetylation during early embryogenesis.

#### Supplementary Tables

**Supplementary Table 1.** Gene signatures used in this work.

**Supplementary Table 2.** Oligonucleotides used to amplify the 3'-UTR of the indicated mouse genes.

**Supplementary Table 3.** Oligonucleotides used for mutagenesis of the miR-203 binding sequences in the 3'-UTR of the indicated mouse genes (binding sites are numbered when more than one is predicted in each 3'UTR).

#### Supplementary References

### Supplementary Figures

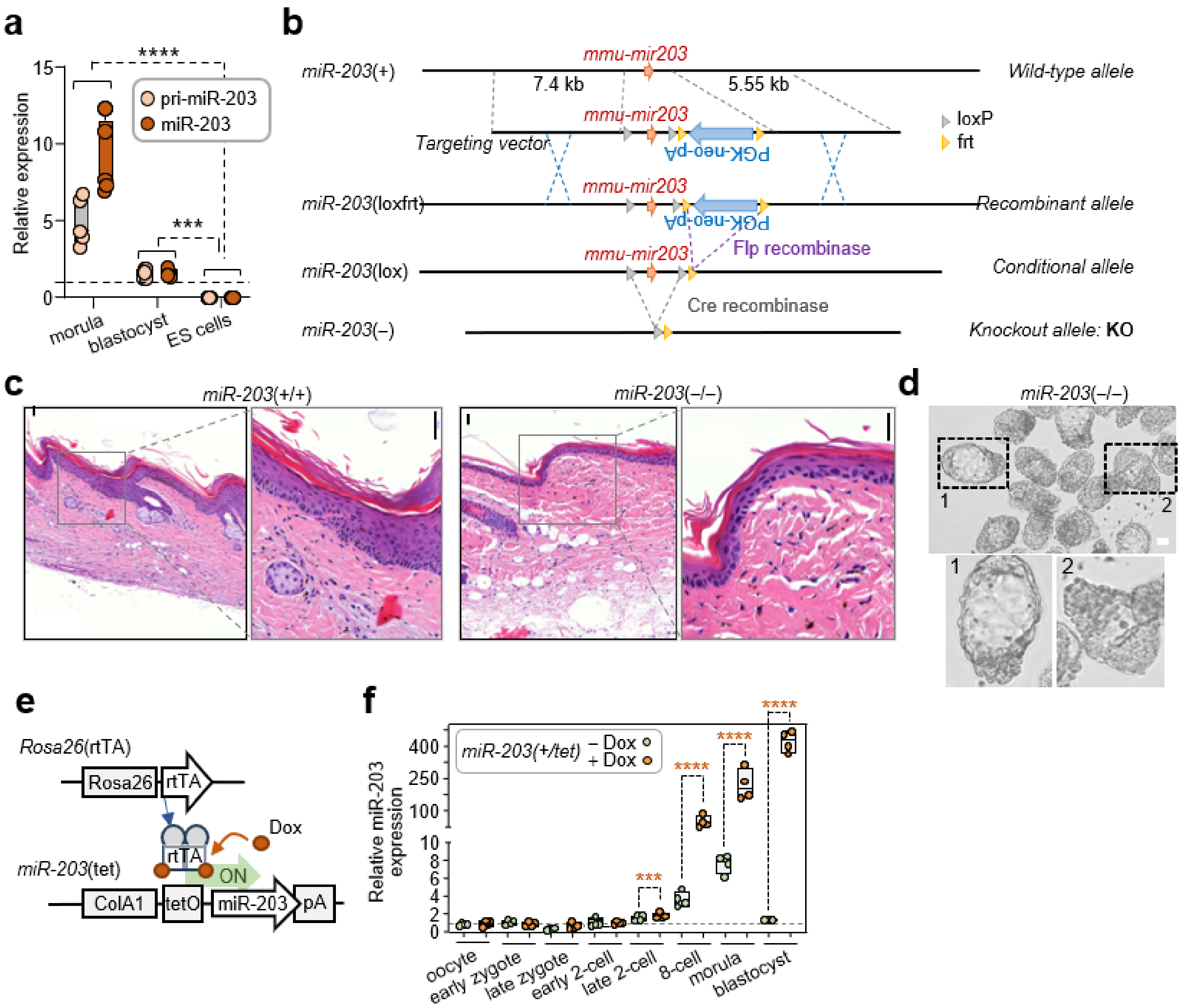


**Supplementary Figure 1. miR-203 expression levels in vivo and generation of miR-203 mutant alleles. a**, miR-203 / pri-miR-203 expression relative to miR-16 / pri-miR-16 in morula (E2.75), blastocyst (E3.5) or established ES cells derived from the miR-203-inducible knock-in model. Each data point represents 25 embryos of the same female donor, or of a different clone of ES cells. Note the absence of pri-miR-203 and miR-203 in ES cell cultures. **b**, Schematic representation of the gene editing strategy for the generation of the *miR-203* conditional knockout mouse model. Note that treatment with tamoxifen promotes the conditional excision of the *mmu-miR-203* gene *miR-203* (lox) to a *mmu-miR-203* (–) knockout allele. **c**, Histological staining by Haematoxylin and Eosin of the skin of adult mice of the indicated genotypes. Scale bar: 100 µm. **d**, Brightfield micrographs of E4.5 *miR-203*(–/–) periimplantation blastocysts in more incipient (1) and advanced (2) developmental stages. Scale bar: 50 µm. **e**, Schematic representation of the miR-203-inducible knock-in model, [*ColA1*(miR-203/miR-203); *Rosa26*(rtTA/rtTA)], in which the reverse tetracycline transactivator is expressed from the *Rosa26* locus, whereas miR-203 is driven by the tetracycline operator downstream of the *ColA1* locus in the presence of doxycycline (Dox). **f**, Expression of *miR-203* in *miR-203*(+/tet) in the absence (green) or presence (orange) of Dox, compared to miR-16 levels in the indicated preimplantation stages, after treatment the mother with Dox when pug was detected. Each data point represents 15-20 embryos of the stated developmental stage from the same female donor (n=~80 total embryos/experimental group). Student t-test with Welsh correction. ***, P<0.001; ****, P<0.0001.

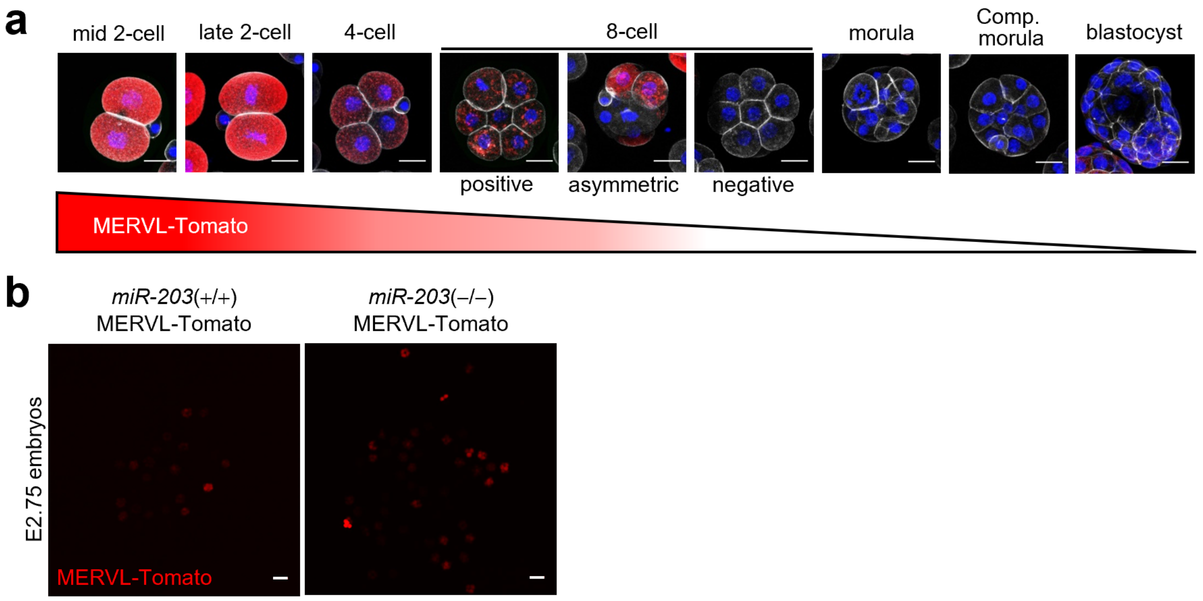


**Supplementary Figure 2. The eE-Reporter mouse registers the activity of MERVL endogenous retroviral elements associated with the totipotent state during early mouse embryogenesis. a**, Confocal micrographs of preimplantational embryos of the indicated stages. Note the paulatino loss of the MERVL-tomato signal (red) throughout the 8-cell stage. 4′,6-diamidino-2-phenylindole (DAPI) is shown in blue. Scale bars: 25 μm. **b**, Confocal micrographs of *miR-203* wildtype and knockout embryos showing MERVL-Tomato signal (red). Scale bars: 100 μm.

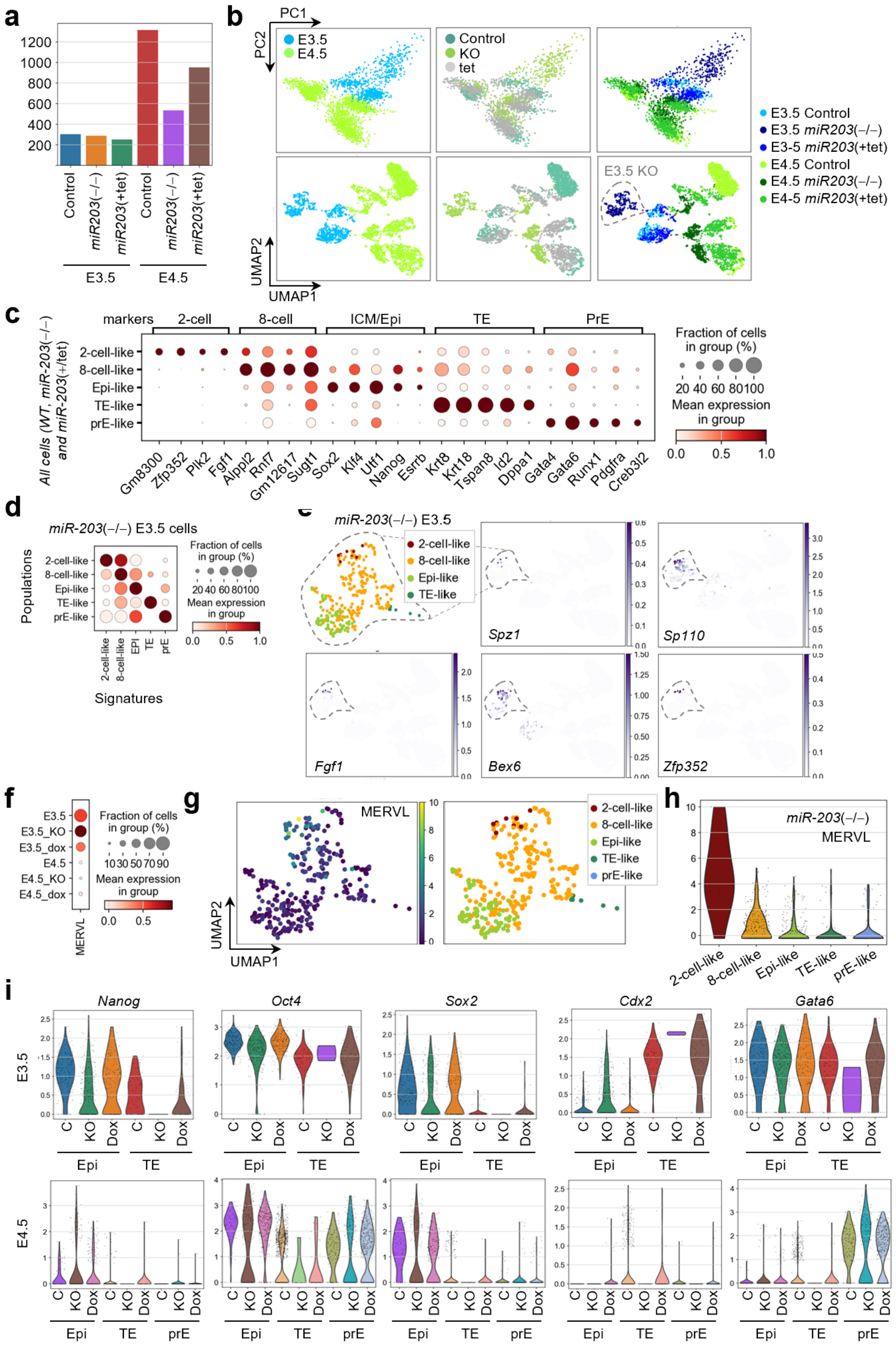


**Supplementary Figure 3. Analysis of cell lineages in miR-203-mutant embryos. a**, Cell counts for each experimental condition subjected to scRNA-seq analysis. **b**, PCA and UMAP plots for cell clusters colored according to developmental stage (left plots), genotype (middle) or stage and genotype (right). **c**, Dot plot representation of the expression of prototypical genes corresponding to the 2-cell, 8-cell, ICM/epiblast, trophectoderm (TE), and primitive endoderm (PrE) identities across the different cell populations segregated in the UMAP plot in **b**. **d**, Dot plot displaying the correlation between the expression of the indicated gene signatures for the different populations contained in the *miR-203*(–/–) E3.5 cell cluster. **e**, Representation of five prototypical genes contained in the 2-cell stage gene signature projected in the UMAP cluster of the *miR-203*(–/–) E3.5 cell cluster. **f**, UMAP plot depicting MERVL expression in the different cell clusters analyzed. Expression bar represents MERVL expression. Note the expression of MERVL in cells contained in the *miR-203*(–/–) E3.5 cell cluster concomitant with the expression of 2-cell genes depicted in **e**. **g**, Dot plot depicting the average expression of MERVL per cell subcluster contained in the *miR-203*(–/–) E3.5 cell cluster.

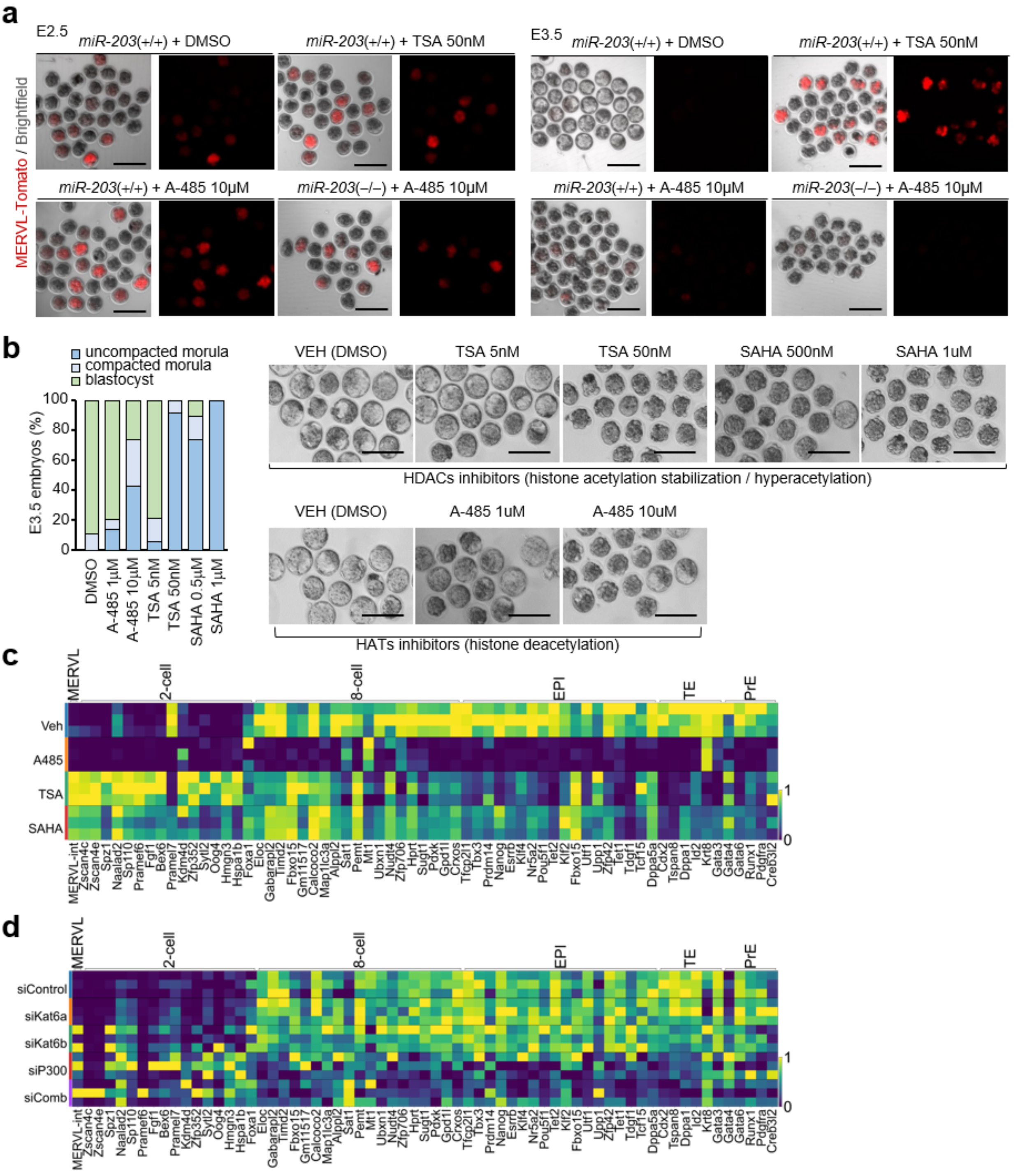


**Supplementary Figure 4**. **Pharmacological modulation histone acetylation during early embryogenesis. a.** Fluorescence and brightfield micrographs of live embryos of the indicated genotypes cultured in vitro since their extraction at E2.5 or E3.5 with the appropriate pharmacological treatments during 6h (E2.5) and 24h (E3.5). Note the presence of MERVL-tomato positive embryos upon treatment with a low dose of the HDAC inhibitor TSA during 24h, mimicking the effect of the loss miR-203, and the induced loss of MERVL-tomato signal upon the treatment with the P300 HAT inhibitor A-485 in *miR-203*(–/–) embryos. Scale bars: 100 µm. **b**, Quantification of the percentage of embryos in each embryonic stage after treatment from E2.75 to E3.5 in vitro with two HAT inhibitors (A-485 for P300 and WM-1119 for KAT6A/B acetyltransferases) and two HDAC inhibitors (trichostatin A: TSA, and Vorinostat: SAHA) at the indicated doses. Right panels show brightfield micrographs of early mouse embryos treated with HAT and HDAC inhibitors at the indicated doses at E2.75 during 24h in vitro. Scale bars: 100 µm.

### Supplementary Tables

**Supplementary Table 1.** Gene signatures used in this work.

| Signatures | Description | Source |
|---|---|---|
| 2-cell | *Zscan4c, Zscan4e, Spz1, Naalad2, Sp110, Pramef6, Fgf1, Bex6, Pramel7, Kdm4d, Zfp352, Sytl2, Oog4, Hmgn3, Hspa1b, Foxa1* | (Rodriguez-Terrones *et al*, 2018) |
| 8-cell | *Dppa3, Map1lc3b, Eloc, Gabarapl2, Timd2, Fbxo15, Gm11517, Calcoco2, Map1lc3a, Alppl2, Sat1, Pemt, Mt1, Ubxn1, Nudt4, Zfp706, Hprt, Sugt1, Pdxk, Gpd1l, Crxos, Ptma, Gm12617, Sumo2, Isyna1, Npm1, Bhmt, Rnf7, Obox6, Mt-rnr2, Eif2s2, Dnaja1, H3f3a, Cks2, Cited1, 2310040g24rik, Ube2c, Cd63, Pttg1ip, Timm17a, Mpc2, Gcsh, Mkrn1, Srp9, Sfn, Tomm5, Dnajb6, Timm23, Gulo, Alg13* | Signature obtained from E2.5 wild-type embryos (unpublished data) |
| Epiblast | *Tfcp2l1, Tbx3, Prdm14, Nanog, Esrrb, Klf4, Nr5a2, Pou5f1, Sox2, Nr0b1, Tet2, Klf2, Fbxo15, Utf1, Upp1, Zfp42, Tet1, Tdgf1, Tcf15, Dppa5a* | (Neagu *et al*, 2020) |
| Trophoectoderm | *Cdx2, Tspan8, Dppa1, Id2, Krt8, Gata3* | (Guo *et al*, 2021) |
| Primitive Endoderm | *Gata4, Gata6, Runx1, Pdgfra, Creb3l2* | (Guo *et al.*, 2021) |

**Supplementary Table 2.** Oligonucleotides used to amplify the 3'-UTR of the indicated mouse genes.

| **Name** | **Forward (5'-3')** | **Reverse (5'-3')** |
|---|---|---|
| *Arid1a* | CAGCCGTGGGACACCTCCCCT | TTACCAAGATTTAATGTACATTTATTCTTAACAC |
| *Arid2* | TGAATCCTACCCCACTGACACAGTG | TAGGCAGTTATAAGTGGATACATTTCACGC |
| *Dr1* | CATTCACCACCTGGGCTTTCAAGAA | AAAACACATCAGAATTTTATTGCTTGAAGA |
| *Ep300* | AGACACCTTGTAGTATTTTGGAAGC | TCAGGTGTCTGTCTCACACAGTTTA |
| *Kat6a* | ATTATACCAGTTGGTATGTGCAGTC | TCCAAAATGGCTCAACATAATTTATTTTTTTATGTT |
| *Kat6b* | ACAGCCTGGCAGCCCACCCA | AAAAGAAAAAACCAAACTCACTTTATTTAAAGACAA |
| *Kmt2c* | ATGCATTCCTTGCTATCTCACCGGG | TATAGATGTAAAATATTTTATTTGTAAATGGTGATCT |
| *Smarcd1* | GGCCTCTGTGGCCCTAGCCT | TGCTCTTTAGAAGTCTTTTATTAAAACTGGGTGA |

**Supplementary Table 3.** Oligonucleotides used for mutagenesis of the miR-203 binding sequences in the 3'-UTR of the indicated mouse genes (binding sites are numbered when more than one is predicted in each 3'UTR)

| **Name** | **Forward (5'-3')** | **Reverse (5'-3')** |
|---|---|---|
| *Arid1a* | CAGTCCTTGCATCAACGGGATGCCACCTAT AATAACTGTTTTTAATGGTTAAAAAAAA | TTTTTTTTAACCATTAAAAACAGTTATTATAGGTG GCATCCCGTTGATGCAAGGACTG |
| *Arid2* | CAGTGGGGTCTCAAAGTCAAATACCTATAA CATACTGTTACTGAAGAAAGCAC | GTGCTTTCTTCAGTAACAGTATGTTATAGGTATT TGACTTTGAGACCCCACTG |
| *Dr1* | ATAATTTATCTTGAACTACACTATGCCCTATA AGAGACTGGCTAATCTTGAAGATTGTC | GACAATCTTCAAGATTAGCCAGTCTCTTATAGG GCATAGTGTAGTTCAAGATAAATTAT |
| *Ep300* | CTCACTTTATGAAAGAATTTAAATAAAAAAAA AGAACCTATATAAAACCAGAGGACAAAAGG GGTTAATGTT | AACATTAACCCCTTTTGTCCTCTGGTTTTATATA GGTTCTTTTTTTTTATTTAAATTCTTTCATAAAGT GAG |
| *Kat6a* | CTTGTTAGTGACTTTGATGCCTTTTAAAATG AGAGCTTTTTCCTATAATTCCATCTTTAAAAT TTTTTATCTT | AAGATAAAAAATTTTAAAGATGGAATTATAGGAA AAAGCTCTCATTTTAAAAGGCATCAAAGTCACT AACAAG |
| *Kat6b-1* | CTGTCTACTCCATGGAAATGCCTTTAGCCTA TAAATTACTGTATATTTTGTTTAAGGTGAC | GTCACCTTAAACAAAATATACAGTAATTTATAGG CTAAAGGCATTTCCATGGAGTAGACAG |
| *Kat6b-2* | AGTGTTTATCTGTCAGGTTTTGAAGCCTATA ATATATGTCCAAATACTTGGCAGGAT | ATCCTGCCAAGTATTTGGACATATATTATAGGCT TCAAAACCTGACAGATAAACACT |

| | | |
|---|---|---|
| *Kat6b-3* | TGAGCTGTGTCATGTGTCACCTATAAGAAC ACATACACGATGGGG | CCCCATCGTGTATGTGTTCTTATAGGTGACACA TGACACAGCTCA |
| *Kmt2c* | TAATTTTAAAATGTGTTTGTATGAACTTGTTT GTTTACCTATATTTAATAAAAAAAAAACCACAC TGTTTTGTGTTTGCTTG | CAAGCAAACACAAAACAGTGTGGTTTTTTTTTA TTAAATATAGGTAAACAAACAAGTTCATACAAAC ACATTTTAAAATTA |
| *Smarcd1* | GTCACAATGAAGAGGGTGTCACCTATATGT CTCACAGTCACCTGTTATC | GATAACAGGTGACTGTGAGACATATAGGTGACA CCCTCTTCATTGTGAC |
